# Regulation potential of transcribed simple repeated sequences in developing neurons

**DOI:** 10.1101/2023.09.04.556210

**Authors:** Tek Hong Chung, Anna Zhuravskaya, Eugene V. Makeyev

## Abstract

Simple repeated sequences (SRSs), defined as tandem iterations of microsatellite- to satellite-sized DNA units, occupy a substantial part of the human genome. Some of these elements are known to be transcribed in the context of repeat expansion disorders. Mounting evidence suggests that the transcription of SRSs may also contribute to normal cellular functions. Here, we used genome-wide bioinformatics approaches to systematically examine SRS transcriptional activity in cells undergoing neuronal differentiation. We identified thousands of long noncoding RNAs containing >200-nucleotide-long SRSs (SRS-lncRNAs), with hundreds of these transcripts significantly upregulated in the neural lineage. We show that SRS-lncRNAs often originate from telomere-proximal regions and that they have a strong potential to form multivalent contacts with a wide range of RNA-binding proteins. Our analyses also uncovered a cluster of neurally upregulated SRS-lncRNAs encoded in a centromere-proximal part of chromosome 9, which underwent an evolutionarily recent segmental duplication. Using a newly established in vitro system for rapid neuronal differentiation of induced pluripotent stem cells, we demonstrate that at least some of the bioinformatically predicted SRS-lncRNAs, including those encoded in the segmentally duplicated part of chromosome 9, indeed increase their expression in developing neurons to readily detectable levels. These data suggest that many SRSs may be expressed in a cell type and developmental stage-specific manner, providing a valuable resource for further studies focused on the functional consequences of SRS-lncRNAs in the normal development of the human brain.

## Introduction

Different types of repeats occupy >50% of the ∼3 billion nucleotide-long human genome, while only ∼1.5% of its capacity is used to encode proteins (Lander et al. 2001; Nurk et al. 2022). The repeated sequences have been traditionally classified as interspersed and tandem repeats, based on their relative positions and the mechanisms driving their expansion. Interspersed repeats often arise through the “selfish” propagation of transposable and retrotransposable elements and their associated sequences(Goodier and Kazazian 2008; Gorbunova et al. 2021). Tandem repeats range from duplications of gene- or gene cluster-sized units to head-to-tail iterations of shorter DNA sequences. Gene duplications can enhance protein activity by increasing the protein production rate or contribute to the processes of evolutionary innovation and speciation (Kent et al. 2003; Lynch and Conery 2000; Nei and Rooney 2005). A notable example of tandem duplication of non-protein-coding genes is provided by the arrays of 47S/45S ribosomal RNA (rRNA) repeats, which are required to sustain the high levels of ribosome biogenesis in the cell (McStay 2023; Nemeth and Grummt 2018).

The biological functions of shorter tandem repeats are less well understood. Depending on the length of their repeated units, this group is often classified into microsatellites (≤9-bp units), minisatellites (10-60-bp units), and satellites (>60-bp units)(Wright and Todd 2023). These boundaries may differ depending on the study and the organism, and alternative terms such as short tandem repeats and simple sequence repeats are often used as synonyms for microsatellites. To avoid confusion, we will collectively refer to all tandem repeats with microsatellite-, minisatellite-, or satellite-sized units as “simple repeated sequences” (SRSs).

Defined in this way, SRSs accounts for ∼7.5% of the total human genome length (Nurk et al. 2022). These repeats, particularly satellites, tend to be enriched in the transcriptionally repressed heterochromatin and associated with genome maintenance functions (Altemose 2022; Ugarkovic et al. 2022). However, at least some SRSs are known to be expressed, giving rise to biologically active transcripts (Ninomiya and Hirose 2020; Trigiante et al. 2021). Since individual SRSs contain multiple repeated units, transcripts containing these sequences can form multivalent contacts with cognate proteins or nucleic acid sequences. This in turn may allow SRS-containing transcripts to act as regulators of cellular RNA metabolism or/and contribute to the assembly of large ribonucleoprotein complexes and membraneless compartments.

Many SRSs are genetically unstable and frequently increase or decrease the number of repeated units as a result of replication or/and recombination errors. The genetic expansion of microsatellites in transcribed genomic regions is known to give rise to toxic RNAs in several neuromuscular and neurodegenerative disorders (Baud et al. 2022; Ciesiolka et al. 2017; Fujino and Nagai 2022; Schwartz et al. 2021). For example, the type 1 myotonic dystrophy (DM1) is caused by the expansion of the CTG trinucleotide repeat in the 3’ untranslated region (3’UTR) of DMPK gene (Meola and Cardani 2015; Sznajder and Swanson 2019; Yum et al. 2017). The aberrant DMPK transcripts containing >50 and, occasionally, up to thousands of CUG units can form a hairpin structure stabilized by the G-C base-pairing.

Furthermore, expanded CUGs comprise numerous YGCY motifs (CUGCUG) recognized by the muscleblind-like (MBNL) RBPs (Mankodi et al. 2001; Miller et al. 2000). This allows for the multivalent binding of MBNL proteins to the mutant DMPK transcripts and the assembly of pathological ribonucleoprotein foci in the nucleus. The sequestration of MBNL proteins and potentially other RBPs results in dysregulation of pre-mRNA splicing and cleavage/polyadenylation, as well as mRNA stability (Goodwin et al. 2015; Jiang et al. 2004; Masuda et al. 2012).

Pathological RNA foci and MBNL sequestration are also observed in the type 2 myotonic dystrophy (DM2), which is caused by the expansion of the CCTG repeats in intron 1 of the CNBP/ZNF9 gene (Liquori et al. 2001; Mankodi et al. 2001; Sznajder and Swanson 2019). Other repeat expansion disorders have been shown to involve the production of toxic RNAs interacting with multiple copies of different RBPs. For example, the expansion of the CAG-repeat in Huntington’s disease (HD; (Nalavade et al. 2013), the GGGGCC-repeat in amyotrophic lateral sclerosis and frontotemporal dementia (ALS and FTD; (Balendra and Isaacs 2018), and the CGG-repeat in Fragile X-Associated Tremor/Ataxia Syndrome (FXTAS; (Cid-Samper et al. 2018; Sellier et al. 2010), have been shown to promote the formation of RNA foci and sequester different protein factors.

The biomedical relevance of SRSs is not limited to repeat expansion disorders. We have previously identified a UC repeat-enriched long noncoding lncRNA (lncRNA), PNCTR, produced by RNA polymerase I (Pol I) from an intergenic spacer separating tandem copies of 47S rRNA genes (Yap et al. 2018). We showed that PNCTR is often upregulated in transformed cells, driving the assembly of the cancer-specific perinucleolar compartment (PNC). The UC-rich sequence elements within PNCTR enable multivalent binding of the RBP PTBP1, sequestering it within the PNC. This in turn antagonizes the splicing regulation function of PTBP1 and promotes cancer cell survival (Yap et al. 2018). Using a proximity labeling approach, we have recently shown that PNCTR and the PNC are associated with a host of additional proteins involved in nucleic acid metabolism (Yap et al. 2022a, b).

SRS transcription can also contribute to normal cellular and organismal processes. A classic example is provided by the lncRNA XIST, which plays a central role in X chromosome inactivation (XCI) in female mammalian cells (Patrat et al. 2020). XCI is essential for the dosage compensation of X chromosome-encoded genes, as females have two X chromosomes, while males have only one. XIST is transcribed by RNA polymerase II (Pol II) from the X chromosome destined to become inactivated (Xi) and facilitates the inactivation process in cis (Patrat et al. 2020).

XIST RNA contains several conserved SRS-like elements, referred to, in the 5’ to 3’ order, as A-, F-, B-, C-, D-, and E-repeats. The F-, B-, C-, and D-repeats contribute to the XIST function by recruiting the RBP hnRNPU, which is required for XIST RNA localization to and spreading along the Xi (Hasegawa et al. 2010; Kolpa et al. 2016; Lu et al. 2020; Sakaguchi et al. 2016). Interaction of the SHARP/SPEN protein with the 5’-proximal A-repeat facilitates the recruitment of the SMRT activator of the histone deacetylase HDAC3, thus promoting the formation of repressive heterochromatin (Chu et al. 2015; McHugh et al. 2015). The A-repeat also forms intra- and inter-molecular AUCG tetraloops, contributing to the multimerisation of Xist RNA in the XCI context (Duszczyk et al. 2011). The E-repeat is Binding of the CIZ1 protein to the E-repeat is involved in the recruitment and anchoring of XIST RNA to the Xi (Ridings-Figueroa et al. 2017; Sunwoo et al. 2017). Furthermore, direct binding of the RPB HNRNPK to the B-, C- and D-repeats is required for the recruitment of a non-canonical Polycomb repressive complex 1 (Almeida et al. 2017; Chu et al. 2015; Lu et al. 2020; Pintacuda et al. 2017).

Another example of a lncRNA expressed in healthy cells is TERRA. This lncRNA enriched in UUAGGG repeats plays a crucial role in the regulation and maintenance of telomeres, protective structures located at the ends of chromosomes. In mice, TERRA can form large and discrete RNA foci in proliferating cells, especially in cells with shortened telomeres (Deng et al. 2012). In yeast, TERRA aids in the recruitment of telomerase clusters to the telomeres, possibly compensating for the limited processivity of telomerase in this organism (Cusanelli et al. 2013). TERRA may also be involved in the telomerase-independent alternative lengthening of telomeres. The formation of an R-loop between TERRA and telomere DNA mediates homology-directed repair in yeasts and telomerase-negative cancer cells, thereby maintaining the telomere length (Graf et al. 2017; Silva et al. 2021).

TERRA also acts as a scaffold for telomere-associated shelterin proteins. The direct interaction between the G-quadruplex formed by the G-rich TERRA RNA and the telomere repeat factor 2 (TRF2) facilitates the recruitment of TERRA to telomeres and the formation of telomeric heterochromatin (Deng et al. 2009; Mei et al. 2021). This interaction also helps to regulate the DNA damage response (Porro et al. 2014). hnRNP A1, a direct binding partner of TERRA, may additionally promote the binding of another shelterin protein, POT1, to exposed single-stranded DNA at the telomeric end, preventing the activation of an ATR-dependent cell-cycle checkpoint (Flynn et al. 2011). TERRA’s association with the histone 3 lysine 9 methyltransferase SUV39H1 may contribute to maintaining the telomere chromatin structure (Arnoult et al. 2012; Porro et al. 2014). Together, TERRA, shelterin proteins, and heterochromatin protect the telomere end from DNA damage response, and the knockdown of TERRA can lead to telomeric defects, chromosome abnormality, and cell death (Barral and Dejardin 2020; Deng et al. 2009).

Similar to telomeres, centromeric and pericentromeric DNA is enriched in SRSs. Transcripts containing centromeric satellite repeats show a considerable sequence diversity across species but play a conserved role in the assembly of the kinetochore and the centromere passenger complex (CPC; (Leclerc and Kitagawa 2021; Perea-Resa and Blower 2018). One common type of centromeric repeats in human is known as alpha satellites, which are characterized by a consensus sequence unit of 171 bp in length (McNulty and Sullivan 2018). Transcripts containing alpha satellites, and possibly other centromeric SRSs, have been shown to associate with the CPC components Aurora B and INCENP, the kinetochore component CENP-C, and the centromeric histone 3 variant CENP-A (Blower 2016; Ideue et al. 2014; McNulty et al. 2017; Quenet and Dalal 2014; Wong et al. 2007).

Knockdown of alpha satellite RNA downregulates these centromeric proteins, resulting in mitotic defects and cell cycle arrest (McNulty et al. 2017; Quenet and Dalal 2014; Wong et al. 2007). Additionally, alpha satellite RNA can associate with SUV39H1, which deposits the repressive heterochromatin marks and recruits the heterochromatin protein 1α (HP1α; (Johnson et al. 2017). Therefore, transcribed alpha satellites can act as scaffolds, facilitating the recruitment of proteins involved in centromeric and pericentromeric chromatin maintenance and chromosome segregation.

Not all satellite-containing RNAs are expressed constitutively. The transcription of the pericentromeric human satellite III (HSATIII) repeats is induced under heat shock and cytotoxic stress conditions by heat shock factor 1 (HSF1) and bromodomain protein 4 (BRD4; (Hussong et al. 2017). The resultant GGAAT repeat-rich HSATIII transcripts drive the assembly of membraneless nuclear stress bodies (nSBs; (Biamonti and Caceres 2009). Examples of proteins localizing to the nSBs include HSF1, Pol II, Scaffold Attachment Factor B (SAFB), SAFB-Like Transcription Modulator (SLTM), Nuclear Receptor Coactivator 5 (NCOA5), hnRNP proteins M, A1, H1, and serine/arginine-rich (SR) proteins such as SRSF1, SRSF7, and SRSF9 (Aly et al. 2019; Denegri et al. 2001; Jolly et al. 2004; Metz et al. 2004; Weighardt et al. 1999). Another nSB component, CDC-like kinase 1 (CLK1), has been shown to mediate SR protein phosphorylation during the recovery from stress (Ninomiya et al. 2020). Phosphorylation of SRSF9 promotes the intron retention in numerous mRNAs, causing their accumulation in the nucleus and dampening the expression of the corresponding genes (Ninomiya et al. 2020).

The above examples indicate that the normal biological functions of SRS-containing transcripts have been extensively studied in proliferating cells. On the other hand, the effects of abnormally expanded SRSs are better understood in diseases affecting differentiated cells, including neurons. The possible roles of transcribed SRSs in healthy neurons remain largely unexplored, partly due to the challenges associated with analyzing repeat-rich sequences. To address this limitation, we systematically identified SRS-containing lncRNA (SRS-lncRNA) candidates upregulated during neuronal development by mining RNA-sequencing (RNA-seq) data. We further validated our bioinformatics predictions using a newly established system for inducible neuronal differentiation of human induced pluripotent stem cells (iPSCs). Our work provides a valuable resource for further studies focused on the roles of repeat transcription in the normal development and function of the nervous system.

## Methods

### Culturing human induced pluripotent stem cells (iPSCs)

Healthy donor-derived iPSCs (HipSci, cat# HPSI0314i-cuhk_1) were maintained in a humidified incubator at 37°C and 5% CO_2_ using Essential 8 (Thermo Fisher Scientific, cat# A1517001) or Essential 8 flex (Thermo Fisher Scientific, cat# A2858501) media supplemented with 100 units/ml PenStrep (Thermo Fisher Scientific, cat# 15140122). Essential 8 medium was changed every day, while Essential 8 flex was refreshed every other day. Tissue culture (TC) plates used for culturing iPSC (typically Thermo Fisher Scientific cat# 140675) were coated by incubating them with 1 µg/cm^2^ of vitronectin (Thermo Fisher Scientific, cat# A14700) at room temperature for 1 hour. For normal passaging, iPSCs colonies were incubated with 0.5 mM EDTA (Thermo Fisher Scientific, cat# 15575020) in DPBS (no calcium, no magnesium; Thermo Fisher Scientific, cat# 14190094) at room temperature for 4-8 minutes and then gently dissociated by trituration in the growth medium.

### DNA constructs

The pZT-C13-L1 and pZT-C13-R1 constructs encoding the left and right TALENs specific to the *CLYBL* safe-harbor locus were a gift from Jizhong Zou (Addgene plasmid #62196 and #62197; (Cerbini et al. 2015). The *CLYBL*-specific homology directed repair construct pUCM-CLYBL-hNIL was a gift from Michael Ward (Addgene plasmid #105841; (Fernandopulle et al. 2018). We modified the pUCM-CLYBL-hNIL backbone to act as an acceptor locus in the recombination-mediated cassette exchange (RMCE) protocol for the high-efficiency integration of transgenes of interest (Iacovino et al. 2011). We used standard molecular cloning techniques and restriction and modification enzymes from New England Biolabs to substitute the *hNIL* fragment with a *Lox2272*- and *LoxP*-flanked *Cre* recombinase gene linked to the *ΔNeoR* and *PuroR* markers. The map of the resultant pML630 plasmid is provided supplementary Supplementary Data S1. The mouse *Ngn2*-encoding RMCE knock-in plasmid pML156 was generated as described previously (A.Z. and E.V.M., submitted).

### Generating TRE-Ngn2 iPSCs

We prepared TRE-Ngn2 iPSCs expressing an *Ngn2* transgene from a Dox-inducible promoter using a two-step approach previously used for mouse ESCs (Iacovino et al. 2011). In our case, the first step involved knocking in the rtTA-2Lox-Cre cassette encoded by pML630 into the *CLYBL* safe-harbor locus, and the second step, high-efficiency RMCE substituting the *Cre* coding sequence with the pML156-encoded *Ngn2*.

In the first step, a ∼20%-confluent wild-type iPSC culture in a 12-well plate containing 1 ml/well of Essential 8 supplemented with 10 µM ROCK inhibitor (Merck, cat# Y0503) was co-transfected with the pZT-C13-L1 and pZT-C13-R1 plasmids (Cerbini et al. 2015) and the pML630 construct mixed in the 1:1:8 ratio, respectively. We combined 2 µg of the plasmid mixture with 2 µl of Lipofectamine™ Stem transfection reagent (Thermo Fisher Scientific, cat# STEM00008) and 100 µl of Opti-MEM I (Thermo Fisher Scientific, cat# 31985070), as recommended. The resultant transfection mixture was then added drop-wise to the cells. The medium was refreshed on the next day. To select knock-in clones, 0.25 µg/ml puromycin was added to the medium 2 days post transfection and gradually increased to 0.75 µg/ml by day 12 post transfection. Puromycin-resistant colonies were picked 12 days post transfection, expanded, and genotyped by PCR using the MLO3670/MLO3671 and MLO3686/MLO1631 primer pairs (Table S1). We also confirmed that the clones express *Cre* in a Dox-inducible manner by reverse transcription (RT) qPCR with using human *GAPDH* as the as the “housekeeping” control (see Table S1 for primer sequences).

In the second step, the rtTA-2Lox-Cre knock-in cells from the first step were pre-treated overnight with 2 µg/ml doxycycline (Dox; Sigma, cat# D9891) to activate *Cre* expression. The cells were then transfected with a mixture containing 1 µg of pML156, 2 µl of the Lipofectamine™ Stem transfection reagent, and 100 µl of Opti-MEM I, as described above. To select RMCE-positive clones, 25 µg/ml G418/geneticin (Sigma, cat# 10131019) was added to the medium 2 days post transfection and gradually increased to 60 µg/ml by day 12 post transfection. G418-resistent colonies were picked two weeks post transfection, expanded and genotyped by PCR with the MLO1295/MLO1296 primers (Table S1). We also confirmed the Dox-inducible expression of the *Ngn2* transgene by RT-qPCR with the same primer pair and using human *GAPDH* as the as the “housekeeping” control. Three TRE-Ngn2 iPSC clones selected in this manner were used for the Dox-induced differentiation experiments described below.

### Dox-induced neuronal differentiation

We adapted the neurogenin 2-based neuronal differentiation protocol from (Fernandopulle et al. 2018) using plates and coverslips coated for 1 hour at 37°C with Geltrex (Thermo Fisher Scientific, cat# A1413302) diluted to 1% with DMEM/F12 (Thermo Fisher Scientific, cat# 31330038). On differentiation day 0, TRE-Ngn2 iPSCs were dissociated with Accutase (Thermo Fisher Scientific, cat# 00-4555-56) at 37°C for 5 minutes and triturated to obtain a single-cell suspension. Cells were then plated onto Geltrex-coated surfaces at 1.5×10^5^ cells/cm^2^ in the induction medium (DMEM/F12 supplemented with 1× N-2 (Thermo Fisher Scientific, cat# 17502048), 1× non-essential amino acids (Thermo Fisher Scientific, cat# 11140035), 1× L-glutamine (Thermo Fisher Scientific, cat# 25030-024), 10 µM ROCK inhibitor, and 2 µg/ml Dox). The medium was replaced on differentiation days 1 and 2, omitting the ROCK inhibitor.

On day 2, we coated fresh 6-well plates (Thermo Fisher Scientific, cat# 140675) or/and 12-well plates (Corning, cat# 3513) with 18-mm coverslips (VWR, cat# 631-1580) with 100 µg/ml poly-L-ornithine (PLO; Merck, cat# A-004-C) and left the plates at 37°C overnight. On day 3, differentiating cultures were dissociated with Accutase at 37°C for 5 minutes and triturated to form a single-cell suspension. Cells were then plated on the PLO-coated plates or coverslips in the maturation media, which consisted of Neurobasal A (Thermo Fisher Scientific, cat# 10888022), 1× B-27 (Thermo Fisher Scientific, cat# 17504044), 10 µg/ml BDNF (Miltenyi Biotec, cat# 130-093-811), 10 µg/ml NT-3 (Miltenyi Biotec, cat# 130-093-973), and 1 µg/ml laminin (Merck, cat# L2020). Dox was omitted from the medium beginning from day 3 and half-medium changes were carried out twice a week until day 14.

### Immunofluorescence

Cells grown on 18-mm coverslips were washed briefly in PBS, fixed with 4% formaldehyde (Thermo Fisher Scientific, cat# 28908), washed three times with PBS, permeabilized with 0.1% Triton X-100 in PBS for 10 minutes, and washed three times with PBS. Fixed and permeabilized cells were then blocked in 0.5% BSA and 0.2% Tween-20 in PBS (IF blocking buffer) for 30 minutes at room temperature and incubated with anti-Ngn2 (Cell Signaling Technology, cat# 13144; 1:250 dilution) and anti-MAP2 (Biolegend, cat# 822501; 1:2000 dilution) antibodies in the IF blocking buffer at 4°C overnight. The samples were washed once with 0.2% Tween-20 in PBS and twice with PBS, and incubated with Alexa Fluor-conjugated secondary antibodies in IF blocking buffers for 1 hour at room temperature. This was followed by one wash with 0.2% Tween-20 in PBS and two washes with PBS. The coverslips were then counterstained with 0.5 µg/ml DAPI in PBS for 3 minutes and mounted with ProLong Gold antifade reagent (Thermo Fisher Scientific, cat# P36934). Images were taken using a ZEISS Axio Observer Z1 Inverted Microscope equipped with a LD Plan-Neofluar 40x/0.6 Corr Ph 2 M27 objective.

### PCR-based assays

PCR genotyping was carried out using the PCRBIO HS Taq Mix Red kit (PCR Biosystems, cat# PB10.23-02), as recommended by the manufacturer. To perform RT-qPCR assays, total RNAs were extracted with the EZ-10 DNAaway RNA Mini-Preps kit (Bio Basic, cat# BS88136) as recommended, and reverse transcribed in a 20-µl format. Prior to reverse transcription, traces of genomic DNA were removed by treating 600 ng of total RNA with 1 µl RQ1 RNase-Free DNase (Promega, cat# M6101) in 9 µl reaction additionally containing RQ1 DNase buffer and 1 µl of murine RNase inhibitor at 37°C for 45 minutes. RQ1 DNase was inactivated by adding 1 µl of Stop Solution from the RQ1 kit and incubating the solution at 65°C for 10 minutes. The reaction was placed on ice, supplemented with 1 µl of 100 µM random decamer primers, incubated at 70°C for 10 minutes, and returned to ice. The reaction was then supplemented with 1× SuperScript IV reaction buffer, 10 mM DTT, 1 mM each of the four dNTPs, 1 µl murine RNase inhibitor, 1 µl of SuperScript IV Reverse Transcriptase (Thermo Fisher Scientific, cat# 18090200), and nuclease-free water (Thermo Fisher Scientific, cat# AM9939) added to the final volume of 20 µl. In RT-negative controls, reverse transcriptase was substituted with an equal volume of nuclease-free water. cDNA was synthesized by incubating the RT mixtures at 50°C for 40 minutes, followed by a 10-minute 70°C heat inactivation step. The samples were finally diluted 10-fold by adding 180 µl of nuclease-free water and stored at −80°C until needed. Quantitative PCR (qPCR) was performed using qPCRBIO SyGreen Master Mix Lo-ROX (PCR Biosystems, cat# PB20.11-51), 100 nM of each primer and 5 µl of diluted cDNA in 20 µl reactions. Reactions were run on a LightCycler 96 Instrument (Roche). The *YWHAZ* gene was used as the “housekeeping” control. Primers used for PCR genotyping and RT-qPCR are listed in Table S1.

### Bioinformatics analyses

A flow chart summarizing our bioinformatic analyses is shown in Fig. 1A. RNA-seq data for the nuclear and cytoplasmic fractions of human embryonic stem cells (ESCs), neural progenitor cells (NPCs), and differentiation day-14 and day-50 neurons were downloaded from Gene Expression Omnibus (https://www.ncbi.nlm.nih.gov/geo/query/acc.cgi?acc=GSE100007; (Blair et al. 2017). RNA-seq data for H1 human ESCs (doi:10.17989/ENCSR895ZTB and doi:10.17989/ENCSR712GOC) and in-vitro differentiated neurons (doi:10.17989/ENCSR877FRY and doi:10.17989/ENCSR877FRY), astrocytes (doi:10.17989/ENCSR129VBC), endothelial cells (doi:10.17989/ENCSR429EGC), and embryonic endodermal cells (doi:10.17989/ENCSR559HWG) were form Encode (https://www.encodeproject.org/). Quality control and adaptor trimming were performed using Trimmomatic (v0.39) (Bolger et al. 2014) as follows:

\# Paired-end data (Blair et al. 2017)

java -jar trimmomatic-0.39.jar PE -threads <threads> \

<input_1.fastq.gz> <input_2.fastq.gz> \

<paired_output_1.fastq.gz> <unpaired_output_1.fastq.gz> \

<paired_output_2.fastq.gz> <unpaired_output_2.fastq.gz> \

LEADING:10 TRAILING:10 SLIDINGWINDOW:5:20 MINLEN:86 \

ILLUMINACLIP:<adaptor.fa>:2:30:10:2:true

#Single-end data (Encode)

java -jar trimmomatic-0.39.jar SE -threads <threads> \

<input.fastq.gz> \

<output.fastq.gz> \

LEADING:10 TRAILING:10 SLIDINGWINDOW:5:20 MINLEN:86 \

ILLUMINACLIP:<adaptor.fa>:2:30:10

**Figure 1.**
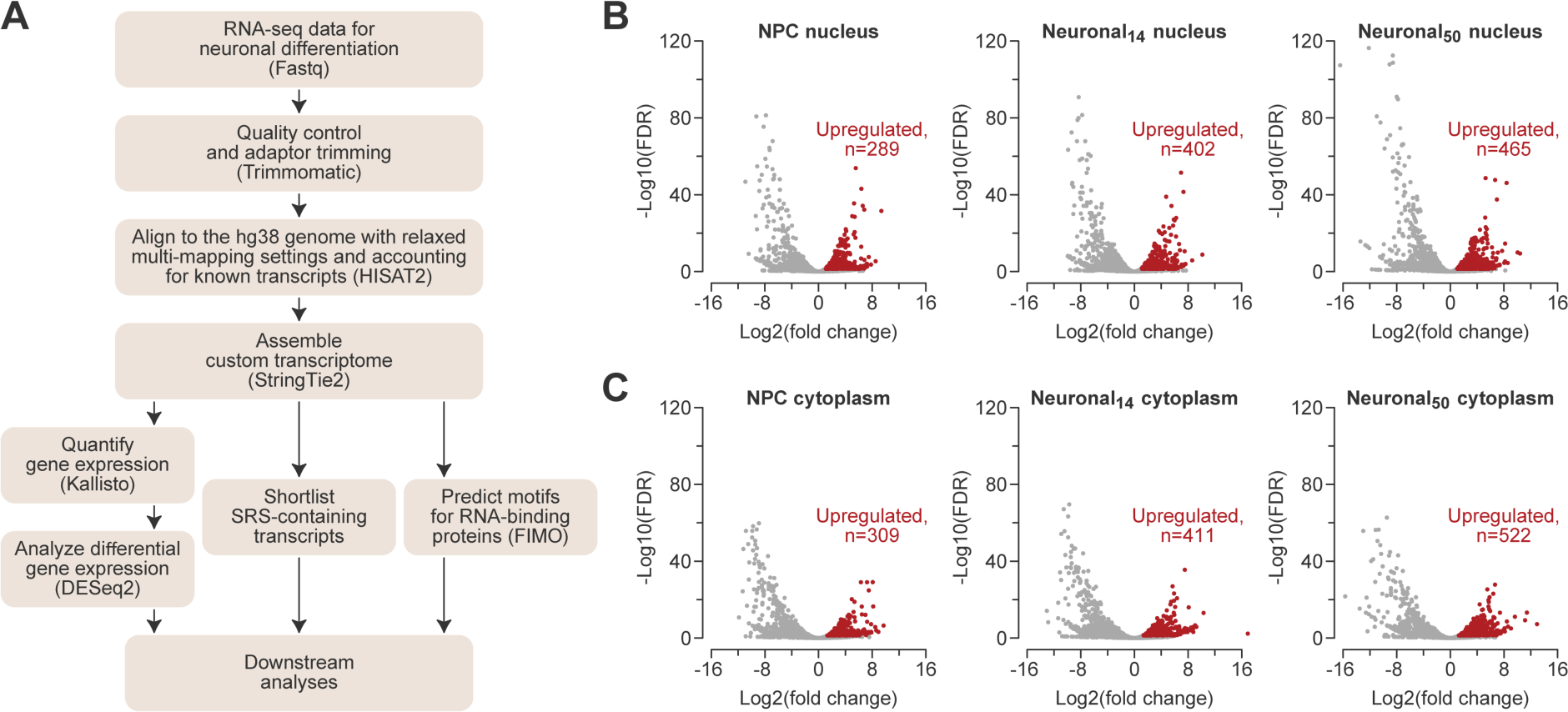
Systematic discovery of SRS-lncRNAs expressed during neuronal differentiation. **(A)** The pipeline used to identify regulated SRS-containing transcripts. **(B-C)** Volcano plots for SRS-lncRNA expression changes in (B) the nuclear and (C) the cytoplasmic fractions at the NPC and day-14 (Neuronal_14_) and day-50 (Neuronal_50_) neuronal differentiation stages, compared to ESC. Significantly upregulated SRS-lncRNA are highlighted in red.

Trimmed FASTQ files were aligned to the GRCh38 genome assembly using HISAT2 (v2.2.1) (Pertea et al. 2016), allowing for RNA-seq read multi-mapping.

#List of known splice sites

python hisat2_extract_splice_sites.py <Gencode V32 annotation GTF file> >

<list_of_known_splice_sites.txt>

#Paired-end data

hisat2 -p <threads> -k 100 --fr --rna-strandness RF --dta-cufflinks --no-unal --no-mixed \

--no-discordant --known-splicesite-infile <list_of_known_splice_sites.txt> \

-x <hg38 genome> \

-1 <trimmomatic paired_output_1.fastq.gz> \

-2 <trimmomatic paired_output_2.fastq.gz> \

-S <output.sam>

#Single-end data

hisat2 -p <threads> -k 100 --rna-strandness R --dta-cufflinks --no-unal \

--no-discordant --known-splicesite-infile <list_of_known_splice_sites.txt> \

-x <hg38 genome> \

-U <trimmomatic output.fastq.gz> \

-S <output.sam>

After converting the output SAM files into indexed BAM files using Samtools (v1.11) (Li et al. 2009), sample-specific transcriptomes were assembled using StringTie2 (v2.1.4) (Pertea et al. 2016) and then merged by Cuffmerge (Trapnell et al. 2010).

stringtie -p <threads> --rf -m 200 -M 1 -c 1 -s 2 -f 0.05 \

-G <Gencode V32 annotation GTF file> -o <output.gtf> <input.bam>

cuffmerge -p <threads> --min-isoform-fraction 0.05 -o <output directory> \

-g <Gencode V32 annotation GTF file> <list_of_gtfs.txt>

The transcripts were quantified using Kallisto (v0.44.0) (Bray et al. 2016) as follows:

#Paired-end data

kallisto quant -i <transcriptome index> --bias --single-overhang --rf-stranded -t <threads> \

-g <cuffmerge_output.gtf> \

-c <chromosome_length.txt> \

-o <output directory> \

<trimmomatic paired_output_1.fastq.gz> <trimmomatic paired_output_2.fastq.gz>

#Single-end data

kallisto quant -i <transcriptome index> --bias --single-overhang --rf-stranded --single \

-l 200 -s 50 -t <threads> \

-g <cuffmerge_output.gtf> \

-c <chromosome_length.txt> \

-o <output directory> \

<trimmomatic output.fastq.gz>

Differential gene expression analyses were performed in DESeq2 using Wald’s test (Love et al. 2014). Genes with log_2_(fold change)>1 and BH-adjusted *P*-value (FDR) <0.05 were considered significantly upregulated, whereas genes with log_2_(fold Change) <-1 and FDR<0.05 were considered downregulated. Low-abundance transcripts expressed at <0.1 transcripts per million (TPM) in the nuclear fraction in all samples were excluded from subsequent analyses.

The transcript_type tag “protein_coding” from the Cuffmerge GTF file output was used to define transcripts and their corresponding genes as protein-coding. The remaining transcripts/genes were considered non-coding. To shortlist SRS-containing transcripts, Simple Repeat track data were downloaded via the UCSC Genome Browser table browser tool selecting the Simple Repeat track for the GRCh38/hg38 genome assembly. Only repeated sequences of >200 bp in length and comprising ≥3 repeated units and were included in downstream analyses. Transcripts consisting entirely of SRS sequences were filtered out because of the inherent difficulty in quantifying their abundance.

To identify RBP-specific sequence motifs, we used FIMO (Bailey et al. 2015) with the RBP motif data from the CISBP-RNA database (Ray et al. 2013):

fimo --text --no-qvalue --norc --thresh 0.001 --motif-pseudo 0.1 \

--max-stored-scores 100000000 \

--bgfile <markov1.b file> \

-oc <output directory> \

<motif file> \

<gene fasta file>

### Statistical analyses

Statistical analyses were carried out in R (v. 4.2.2; (R_Core_Team 2021). Data were compared using two-sided Student’s t-test or Wilcoxon signed-rank test, or one-sided Wilcoxon rank sum test. Multiple pairwise comparisons were adjusted for multiple testing using the Benjamini-Hochberg FDR method. Alternatively, we used one-way ANOVA followed by Tukey’s post-hoc test or the Kruskal-Wallis test followed by Dunn’s post-hoc test, as indicated in the Figures. SRS-lncRNA distributions along the centromeric-to-telomeric were compared using Kolmogorov-Smirnov test. Fisher’s exact test was used to compare categorical data. *P*-values <0.05 were considered statistically significant.

## Results

### Widespread regulation of SRS-lncRNA expression during neuronal differentiation

To investigate the transcriptional status of SRSs encoded in the reference human genome, we examined a publicly available RNA-sequencing (RNA-seq) dataset for human embryonic stem cells (ESCs) undergoing *in vitro* differentiation into forebrain-specific neurons (Blair et al. 2017). We selected this study since it contains high-quality sequencing data for four differentiation stages: ESCs, neural progenitor cells (NPCs), and neurons at two stages of maturation, 14 days (Neuronal_14_) and 50 days post-differentiation from NPCs (Neuronal_50_). Furthermore, we reasoned that the nuclear- and cytoplasmic-fraction data provided by this study should increase the likelihood of identifying compartment-enriched transcripts.

Our RNA-seq analysis pipeline was designed to handle multi-mapping reads and incorporated reference-guided transcriptome assembly, allowing us to include both previously annotated and novel RNAs (Fig. 1A; see Methods for further details). We defined SRS-containing long noncoding RNAs (lncRNAs) as transcripts that lack a known protein-coding sequence and contain >200 nt-long simple repeat(s) (with at least 3 repeated units) sourced from the UCSC Genome Browser database. The choice of the >200-nt cutoff was based on the traditional definition of lncRNAs (Mattick et al. 2023). To mitigate the artifacts of the ambiguous alignment of multi-mapping reads, we excluded newly predicted transcripts composed entirely of SRSs. This uncovered a total of 5430 SRS-lncRNA candidates expressed in the nucleus or/and cytoplasm in at least one of the differentiation stages with the >0.1 TPM cutoff.

To identify differentially expressed SRS-lncRNAs, we compared the NPC, Neuronal_14_ or Neuronal_50_ samples to the corresponding ESC controls (Fig. 1A). This revealed 899 non-redundant candidates that were upregulated either in the nucleus or the cytoplasm (>2-fold, FDR<0.05) in at least one neural sample (i.e. in NPC, Neuronal_14_ or Neuronal_50_; Table S2). Among the upregulated SRS-lncRNAs, 289 SRS-lncRNAs were upregulated in the NPC nuclei and 402 and 465 in the Neuronal_14_ and Neuronal_50_ nuclei, respectively. Similarly, 309 SRS-lncRNAs were upregulated in the NPC cytoplasm and 411 and 522 in the Neuronal_14_ and Neuronal_50_ cytoplasm (Fig. 1B-C). Our pipeline also identified 444 ESC-specific candidates that were consistently downregulated at all the neural stages in at least one compartment (>2-fold, FDR<0.05) and not significantly upregulated in the other compartment (Table S3).

The SRS-lncRNAs shortlisted in this manner included previously characterized transcripts with relevant expression patterns. For instance, the LINC00632 (XLOC_334612) precursor of the brain-enriched circular RNA CDR1as/CiRS-7 containing multiple microRNA miR-7-interacting sequences (Barrett et al. 2017; Hansen et al. 2013; Memczak et al. 2013), was upregulated in NPCs and neurons (Fig. S1A and Table S2). Conversely, the lncRNA CPMER (XLOC_185088) involved in cardiomyocyte differentiation and expressed in embryonic stem cells (Lyu et al. 2022), as well as the p53-regulated pluripotency-specific lncRNA LNCPRESS2 (XLOC_222487; (Jain et al. 2016) were downregulated in neural samples (Fig. S1B-C and Table S3). These examples provided internal controls for the performance of our pipeline. Based on the inspection of UCSC Genome Browser data, many other shortlisted transcripts were either novel (e.g., Fig. S2A-B) or matched known lncRNAs not previously associated with the neural lineage (e.g., Fig. S2C).

Thus, many SRS-containing transcripts may change their expression during normal neuronal differentiation.

### Bioinformatics characterization of SRS-lncRNAs upregulated in neural cells

To gain initial insights into the 899 SRS-lncRNAs upregulated in NPCs or/and neurons, we compared the positions of their loci with those of all SRSs encoded in the genome and all detectably expressed SRS-lncRNAs. Plotting the overall genomic distribution of SRSs along the centromere-to-telomere axis revealed their expected enrichment in the centromere- and telomere-proximal regions (Fig. 2A). In comparison, all detectable SRS-lncRNAs lost the centromere-proximal peak and showed significant enrichment in a broader telomere-proximal region (Fig. 2B).

**Figure 2.**
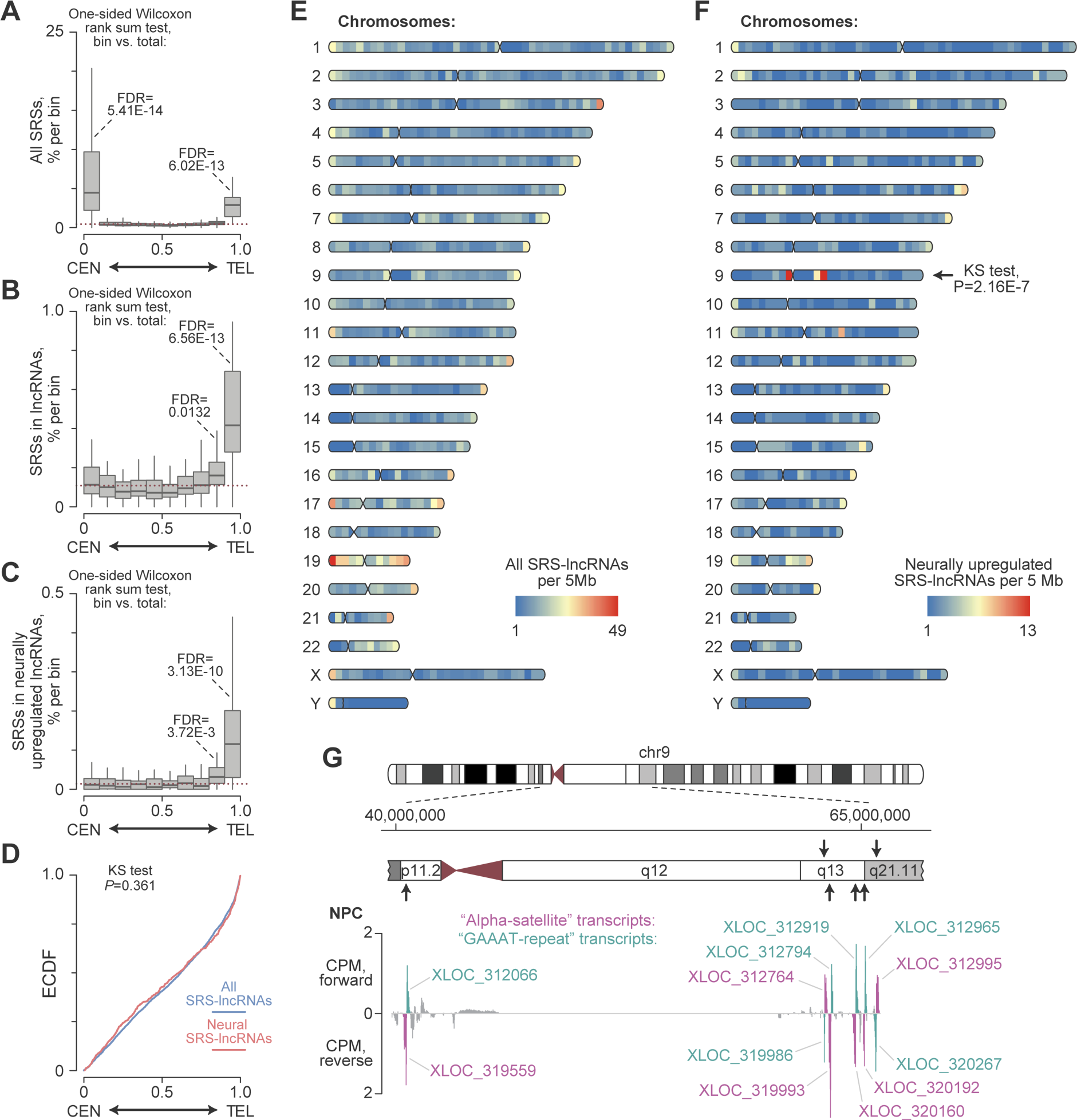
Genomic distribution of SRS-lncRNAs. **(A-C)** Box plots showing the incidence of (A) all genome-encoded SRSs of >200 bp in length and containing ≥3 repeated units, (B) SRSs in all detectable expressed lncRNAs, and (C) SRSs in neurally upregulated lncRNAs along the human chromosome arms separated into 10 equally sized bins, form the middle of the centromere (position 0) to the end of the telomere (position 1). The p-arms of the acrocentric chromosomes 13, 14, 15, 21 and 22 encoding the *47S/45S* rRNA arrays were excluded from this analysis. **(D)** The Kolmogorov-Smirnov (KS) test shows that the distribution of all SRS-lncRNAs (blue) and neural SRS-lncRNAs (red) along the centromere-to-telomere axis do not significantly differ on a genome-wide scale. The data are presented as empirical cumulative distribution function (ECDF) plots. **(E-F)** Karyoplots showing the color-coded density of (E) all detectably expressed SRS-lncRNAs and (F) neurally upregulated SRS-lncRNAs, calculated for individual human chromosomes as the number of loci per a 5-Mb window. The centromere-to-telomere distributions of the neurally upregulated SRS-lncRNAs and all detectably expressed SRS-lncRNAs were compared for individual chromosome arms using the KS test. A significant difference was detected only for chromosome 9 (chr9). The panels were generated using the Rideogram package (https://cran.r-project.org/web/packages/RIdeogram/vignettes/RIdeogram.html) **(G)** A close-up of the centromere-proximal part of chr9 encoding twelve neurally upregulated SRS-lncRNAs organized as six segmentally duplicated bidirectional transcription units (arrows). The panel also displays strand-specific counts-per-million (CPM) normalized coverage plots for the nuclear NPC RNA-seq data. The alpha satellite- and the GAAAT repeat-containing SRS-lncRNAs, which constitute the bidirectional unit, are highlighted in magenta and teal, respectively. The coverage plot on the top represents RNAs transcribed in the forward direction, while the bottom part shows RNAs transcribed in the reverse direction.

The subset of neural SRS-lncRNAs exhibited a similar telomere-biased profile (Fig. 2C-D). However, further examination at the individual chromosome level revealed specific enrichment of neural SRS-lncRNAs in centromere-proximal parts of chr9 comprising evolutionarily recent segmental duplications (Fig. 2E-F; (Bailey et al. 2002; Crosier et al. 2002; Guy et al. 2000). Interestingly, twelve SRS-lncRNAs encoded in this region are organized into six bidirectional transcription units sharing considerable sequence similarity. Within each unit, one SRS-lncRNA typically contains a 10-36 kb-long alpha-satellite sequence, while the SRS-lncRNA transcribed in the opposite direction has a 3-8 kb-long GAAAT-rich microsatellite repeat (Fig. 2G).

Several previously characterized SRS-containing transcripts have been shown to recruit multiple copies of specific RBPs (e.g., (Yap et al. 2018). With this in mind, we conducted a sequence motif analysis and found that 695 out of the 899 neurally upregulated SRS-lncRNAs contain ≥10 putative interaction sites for individual RBPs within their SRS regions (Table S4; see Methods for further details). Importantly, the motif density was significantly higher in the SRS regions compared to the non-SRS parts of the same SRS-lncRNA (Fig. 3A). Furthermore, many SRS-lncRNAs were predicted to engage in multivalent interactions with more than one distinct type of RPBs, with a median of 5 different RBPs (Fig. 3B).

**Figure 3.**
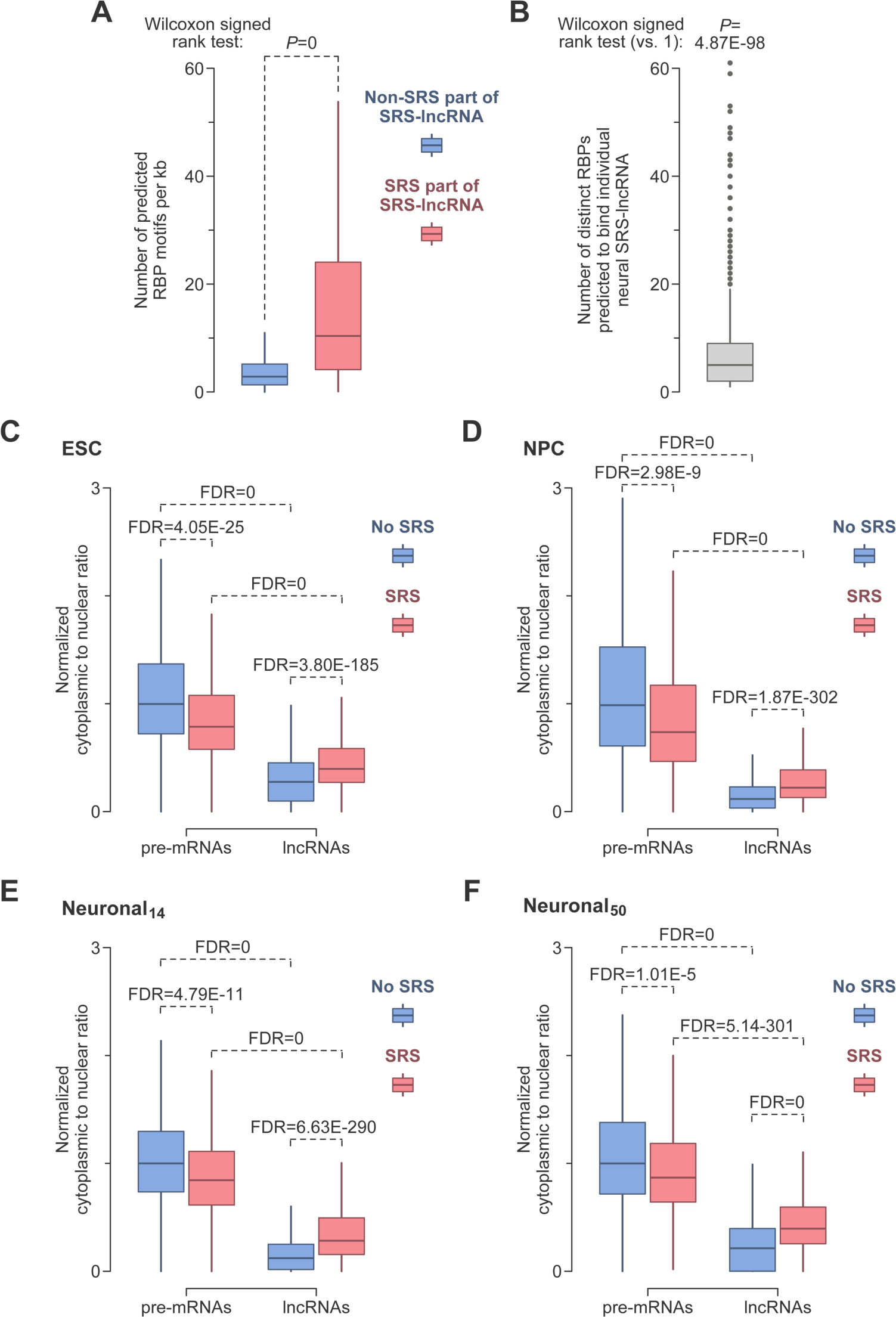
Bioinformatics analyses of SRS-lncRNAs. **(A)** The density of predicted RBP motifs is significantly higher in the SRS parts of SRS-lncRNAs compared to the non-SRS parts of the same transcript. **(B)** Individual SRS-lncRNAs expressed in the neural lineage frequently contain ≥10 interaction motifs for more that one type of RBP. **(C-F)** Non-SRS pre-mRNA median-normalized ratios between transcript abundances in the cytoplasm and the nucleus for the (C) ESC, (D) NPC, (E) Neuronal_14_ and (F) Neuronal_50_ samples. The data were compared by the Kruskal-Wallis test (*P*=0 for all panels) with Dunn’s post-hoc test (*P*-values shown in the panels). The data in (A-F) are presented as box plots, with outliers shown in (B) but not the other panels.

Since many lncRNAs are retained in the nucleus (Guo et al. 2020; Palazzo and Lee 2018; Tong and Yin 2021), we compared the intracellular distribution of SRS-lncRNAs to that of other types of transcripts. As expected, the cytoplasmic-to-nuclear ratio was noticeably lower for SRS-lncRNAs than for primary protein-coding transcript (pre-mRNAs) with or without >200 nt-long SRSs. However, SRS-lncRNAs were relatively more abundant in the cytoplasm compared to lncRNAs lacking qualifying SRSs. Interestingly, the presence of such repeats in pre-mRNAs led to the opposite effect – a decrease in the cytoplasmic-to-nuclear ratio. These effects were observed for transcripts detectably expressed at all four stages of differentiation (Fig. 3C-F; >0.1 TPM in the nuclear fraction at the corresponding stage).

These data reveal the genome distribution SRS-lncRNAs expressed in the neural lineage and suggest that these transcripts have a strong potential for RBP recruitment. Moreover, our analyses are consistent with the possibility that SRSs play a role in regulating the relative abundance of RNA in the nucleus and cytoplasm, in a transcript type-dependent manner.

### Doxycycline-inducible differentiation of human iPSCs into neurons

To validate the SRS-lncRNA regulation patterns, we generated a stable human iPSC line with a doxycycline (Dox) inducible *neurogenin 2* (*Ngn2*) transgene (Fig. S3A). Ectopic expression of Ngn2 is known to trigger efficient neuronal differentiation of proliferating stem cells *in vitro* (Fernandopulle et al. 2018; Lin et al. 2021; Zhang et al. 2013). Inspired by the high-efficiency RMCE-based knock-in technology available for mouse ESCs (Iacovino et al., 2011), we first knocked in an “acceptor” cassette into the *CLYBL* safe-harbor locus of wild-type human iPSCs using the appropriate TALEN and homology-directed repair constructs (Fig. S3A). The cassette contained a puromycin selection marker, a constitutively expressed reverse tetracycline transactivator (rtTA) and a tetracycline/doxycycline responsive element (*TRE*) promoter driving the expression of a *lox2272* and *loxP* site-flanked *Cre* recombinase gene. To enable the subsequent RMCE-dependent integration step, the cassette also encoded a fragment of the G418 resistance gene (*ΔNeoR*) that lacked a promoter and an in-frame translation initiation codon.

After selecting the rtTA-2Lox-Cre line resistant to puromycin (but not G418), we proceeded with the RMCE-dependent knock-in step. The cells were treated with Dox to induce *Cre* expression and transfected with the pML156 “donor” plasmid containing the mouse *Ngn2* along with the *lox2272* and *loxP* sites (Fig. S3A). The pML156 design also precluded transient expression of *Ngn2* that might promote unwanted iPSC differentiation. Cre-mediated RMCE was expected to integrate the *Ngn2* transgene under the *TRE* promoter, replacing the *Cre* gene. Furthermore, the *ΔNeoR* fragment was supposed to be appended with a start coding and the human PGK promoter (*hPGK*), making the resultant TRE-Ngn2 cells G418 resistant. We confirmed *Ngn2* integration by PCR-genotyping G418-resistant clones (Fig. S3B). Moreover, TRE-Ngn2 cultures treated with Dox for 48 hours expressed readily detectable amounts of the Ngn2 protein and the neuronal marker MAP2 (Fig. S3C). Expression of these proteins was not detected in mock-treated cells (Fig. S3C).

To induce neuronal differentiation, we incubated the TRE-Ngn2 cells with Dox for three days and then maintained the cultures until day 14 in a medium supplemented with the neurotrophic factors BDNF and NT-3 (Fig. 4A). RT-qPCR analyses of RNA samples collected on differentiation days 0, 1, 2, 3, 7, and 14 showed that the differentiating cells progressively lost the expression of the pluripotency markers POU5F1 and NANOG (Fig. S3D-E) and gained the expression of an early neuronal marker, NCAM1, beginning from differentiation day 1 (Fig. S3F). The mature neuronal marker SYN1 and the glutamatergic neuronal marker SLC17A6/VGLUT2 were upregulated on differentiation day 7 and continued to be expressed on day 14 (Fig. S3G-H).

**Figure 4.**
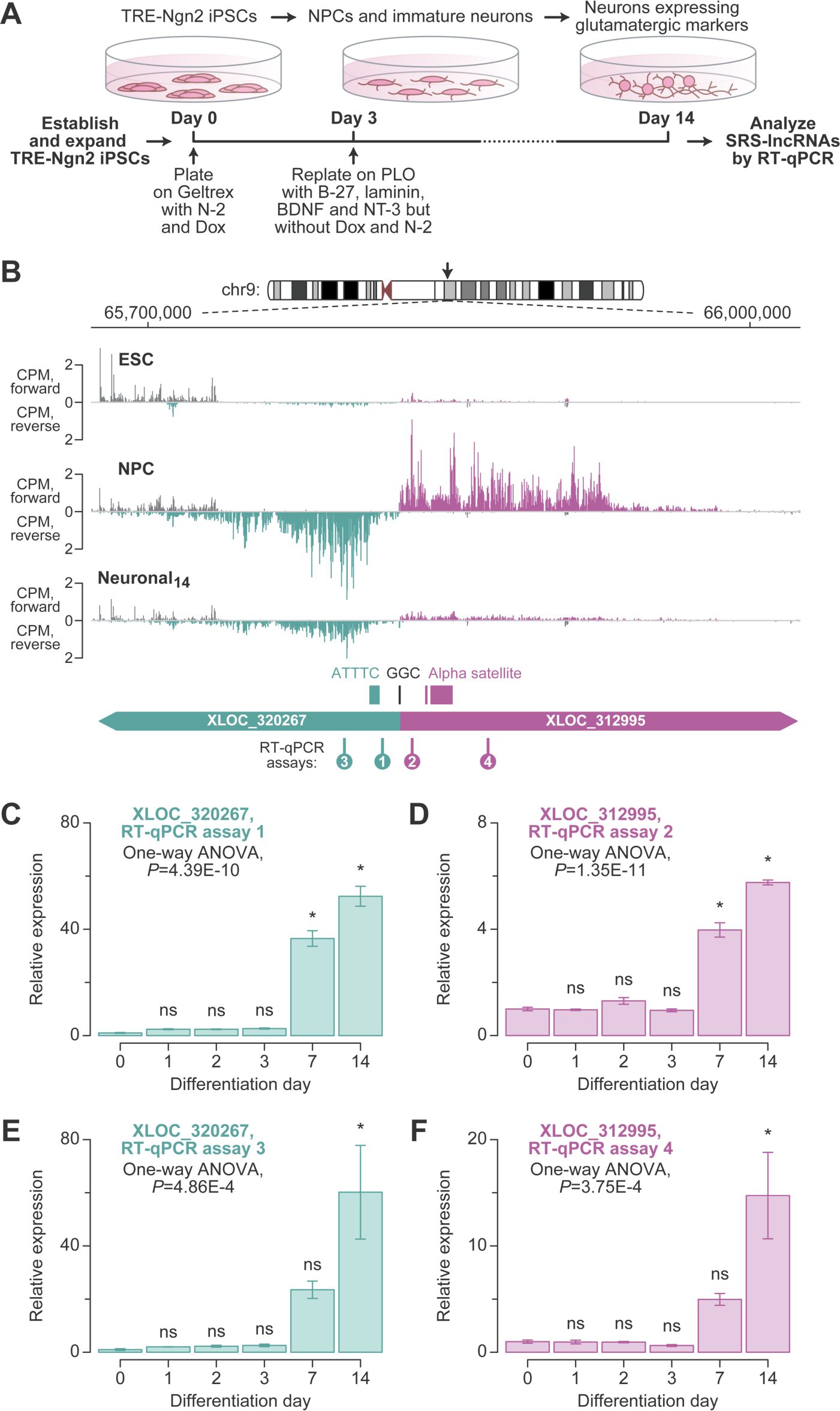
Neuronal differentiation system for SRS-lncRNA validation. **(A)** The protocol for Dox-inducible neuronal differentiation of TRE-Ngn2 iPSCs established in this study (see Methods for further details). **(B)** CPM-normalized RNA-seq coverage plots for the nuclear fraction of ESC, NPC, and Neuronal_14_ samples, illustrating one of the six segmentally duplicated chr9 SRS-lncRNA units. The unit encodes the “alpha-satellite” (XLOC_312995; magenta) and the “GAAAT-repeat” (XLOC_320267; teal) SRS-lncRNAs transcribed in the opposite directions from a common GC-rich (GGC) promoter region. The two SRS-lncRNAs and the corresponding SRSs are annotated at the bottom. Note that the microsatellite sequences are shown for the forward strand. The diagram also shows the positions of the four RT-qPCR amplicons (assays 1-4) analyzed in (C-F). **(C-F)** RT-qPCR assays validating the regulation of (C, E) XLOC_320267 and (D, F) XLOC_312995 expression during neuronal differentiation using primer pairs annealing either upstream (C and D) or downstream (E and F) for the corresponding SRSs. Data were averaged for differentiation experiments carried out using three distinct TRE-Ngn2 iPSC clones ±SEM and compared by one-way ANOVA with Tukey’s post-hoc test. *, Tukey’s *P*<0.05; ns, Tukey’s P≥0.05.

We concluded that Dox-induced TRE-Ngn2 cultures recapitulated the natural dynamics of gene expression in developing neurons.

### Validation of neurally upregulated SRS-lncRNAs

We then utilized the newly established neuronal differentiation system to validate our bioinformatic predictions. TRE-Ngn2 iPSCs were differentiated as outlined in Fig. 4A and analyzed by RT-qPCR on differentiation days 0, 1, 2, 3, 7, and 14, using appropriate primers. We first investigated the clustered SRS-lncRNAs from the segmentally duplicated part of chr9 (Fig. 2G). According to the RNA-seq data, both members of the bidirectionally transcribed unit were expected to be upregulated in NPCs and retain detectable expression in neurons, albeit at a somewhat decreased level (Fig. 4B).

Our RT-qPCR analyses of the TRE-Ngn2 time series with primers annealing upstream of the alpha-satellite and the GAAAT repeats confirmed the upregulation of the corresponding SRS-lncRNAs (e.g., XLOC_312995 and XLOC_320267; see Fig. 2G) during neuronal differentiation (assays 1 and 2; Fig. 4B-D). Similar results were obtained when we repeated the analysis using downstream primers designed against downstream sequences (assays 3 and 4; Fig. 4B, E-F). Of note, RT-qPCR showed a higher expression of the two types of SRS-lncRNAs in day-14 neurons compared to the earlier stages of TRE-Ngn2 differentiation. The apparent difference of this dynamics from Fig. 4B might be due to possible variability between the ESC and the TRE-Ngn2 iPSC differentiation protocols or the fact the RNA-seq data were obtained for the nuclear fraction, while the RT-qPCR was performed for whole-cell RNA samples.

We then carried out longitudinal RT-qPCR analyses for the neurally upregulated SRS-lncRNA candidates XLOC_319631, XLOC_320382, and XLOC_039609/KRTAP5-AS1 (Fig. S2) using SRS-proximal primers. In all 3 cases, the expression significantly increased as a function of development, peaking in day-7-14 neurons (Fig. 5A-C). Conversely, the neurally downregulated SRS-lncRNA candidate XLOC_185088/CPMER (Fig. S1B) decreased its expression in the RT-qPCR time-course, as expected (Fig. 5D).

**Figure 5.**
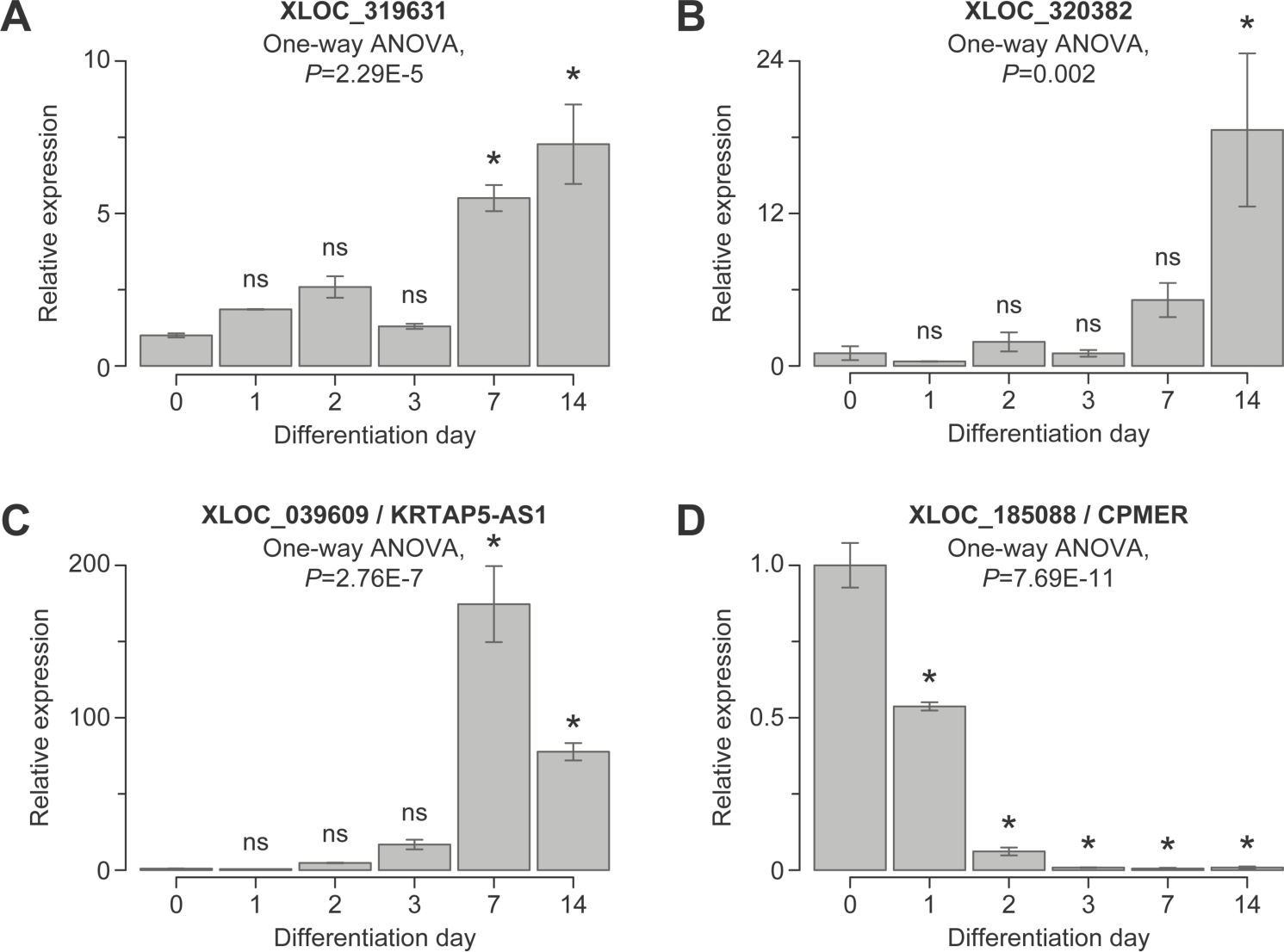
Experimental validation of SRS-lncRNAs. We used RT-qPCR with SRS-proximal primers to validate the expression dynamics of **(A)** XLOC_319631, **(B)** XLOC_320382, and **(C)** XLOC_039609/KRTAP5-AS1, which are predicted to be upregulated during neuronal differentiation, as well as **(D)** XLOC_185088/CPMER, which is predicted to be downregulated. Data were averaged for differentiation experiments carried out using three distinct TRE-Ngn2 iPSC clones ±SEM and compared by one-way ANOVA with Tukey’s post-hoc test. *, Tukey’s *P*<0.05; ns, Tukey’s P≥0.05.

We finally asked if the neural SRS-lncRNAs identified in an ESC differentiation experiment were also upregulated in RNA-seq data available for neurons or/and other types of differentiated cells. To this end, we mined Encode RNA-seq data and shortlisted SRS-lncRNAs upregulated in specific types of human cells compared to ESCs. The Encode list for neurons showed a robust overlap with the neurally upregulated SRS-lncRNAs selected by our bioinformatics pipeline (321 out of 899; Fig. 6A and Table S5). Other cell types, including astroglia, endothelial, and endodermal cells, also shared considerable numbers of common SRS-lncRNAs with the neurally upregulated SRS-lncRNAs (Fig. 6B-C and Table S5), but the overlaps in these cases were smaller compared to Fig. 6A. Notably, 129 out of the 321 overlapping neuronal SRS-lncRNAs were not upregulated in the other cell types (Table S5). This subset included, for instance, the XLOC_039609/KRTAP5-AS1 transcript (Fig. S2) and two out of the six GAAAT repeat-containing transcripts from the chr9 SRS-lncRNA cluster (XLOC_312965 and XLOC_319986; Fig. 2G).

**Figure 6.**
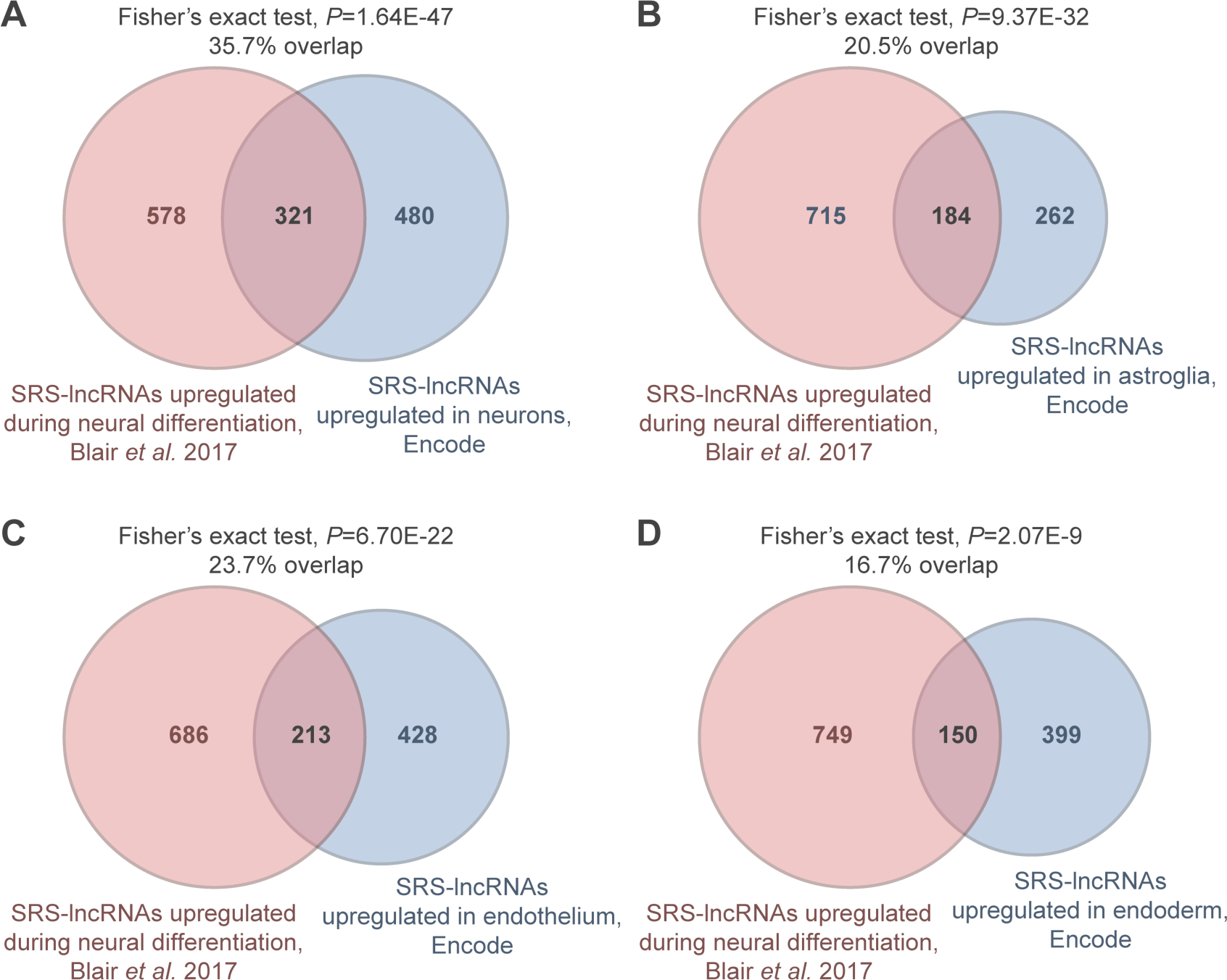
SRS-lncRNA expression across differentiated cell types. Venn diagrams show overlaps between the neurally upregulated SRS-lncRNAs identified by our pipeline using the data from (Blair et al. 2017) and SRS-lncRNAs upregulated in differentiated **(A)** neuronal, **(B)** astroglial, **(C)** endothelial and **(D)** endodermal cells from the Encode database. Note that differentiated neurons show the most robust overlap.

Thus, many SRS-lncRNAs predicted by our pipeline are expressed at readily detectable levels and consistently upregulated in developing neurons.

## Discussion

Our study argues that numerous long noncoding transcripts containing extensive stretches of simple repeated sequences are expressed in genetically normal human pluripotent stem cells and developing neurons (Fig. 1). Many of these SRS-lncRNA originate from telomere-proximal regions, despite the abundance of SRSs in both centromere- and telomere-proximal DNA (Fig. 2). While additional work will be required to understand the molecular basis of this genome-wide bias, possible underlying reasons might include more accessible structure or/and more favorable epigenetic modifications of the telomere-proximal chromatin compared to its centromere-proximal counterpart.

Importantly, a large fraction of the SRS-lncRNAs predicted by our bioinformatics pipeline is significantly upregulated during normal neuronal differentiation, and at least some of these transcripts appear to be specific to neurons (Fig.1, Figs. 4-6, Table S2, and Table S5). Similar to other SRS-lncRNAs, such neural transcripts tend to be encoded in telomere-proximal regions of the genome. An important exception is a broad centromere-proximal region of chr9 that gives rise to a significantly larger number of neural SRS-lncRNAs than expected by chance (Fig. 2E-F). The SRS-lncRNAs encoded in this region are organized as bidirectional pairs of the alpha satellite- and GAAAT repeat-containing transcripts expressed from a common GC-rich promoter (Fig. 2G and Fig. 4B). This part of the genome is known to be segmentally duplicated in primates (Bailey et al. 2002; Crosier et al. 2002; Guy et al. 2000), suggesting an intriguing possibility that it encodes primate-specific functions.

Our RBP motif analyses suggest that the chr9-derived and other SRS-lncRNAs upregulated in NPCs and neurons can recruit multiple copies of specific RBPs via SRS-enriched cognate sequence motifs (Fig. 3A-B and Table S4). For example, the medium numbers of multivalent RBP motifs in the SRS parts of the alpha-satellite and GAAAT-repeat SRS-lncRNA families enriched on chr9 are 46 and 31.5, respectively (Table S4). Validating such interactions and understanding their possible role in the RNA metabolism of developing neurons will be an exciting direction for future studies.

It will be also interesting to follow up on our finding that SRSs are associated with a decrease in the cytoplasmic-to-nuclear ratio for protein-coding transcripts, while having the opposite effect on lncRNAs (Fig. 3C-F). We hypothesize that this difference relates to the abundance of introns in the former group and their paucity in the latter. Indeed, at least some SRSs are known to interfere with intron excision and mRNA export from the nucleus to the cytoplasm (Monteuuis et al. 2019; Sznajder et al. 2018; Yap et al. 2012).

Although SRS-lnRNAs are clearly more nucleus-biased than protein-coding transcripts (Fig. 3C-F), it is tempting to speculate that the increased cytoplasmic abundance of SRS-lnRNAs compared to other lncRNAs is functionally relevant. A previously characterized example of a repeat-containing lncRNA that acts in both the nucleus and the cytoplasm is NORAD (Elguindy and Mendell 2021; Lee et al. 2016; Munschauer et al. 2018; Tichon et al. 2016). This conserved transcript lacks classically defined SRSs, but contains 18 UGURUAUA motifs that may have originated from ancient duplication events. NORAD utilizes these sequences to sequester Pumilio-family RBPs, thereby altering mRNA stability in the cytoplasm.

Another relevant example is provided by competing endogenous RNAs (ceRNAs) that regulate microRNA activity in the cytoplasm (Ala 2020). Interestingly, our analysis identified two known ceRNA candidates: the circular RNA CDR1as/CiRS-7 and the lncRNA XLOC_039609/KRTAP5-AS1 (Fig. S1A and Fig. S2C; (Barrett et al. 2017; Hansen et al. 2013; Memczak et al. 2013; Song et al. 2017). We are currently investigating the possibility that other cytoplasmically abundant SRS-lncRNAs might interact with specific microRNAs.

In conclusion, we have identified multiple instances of simple repeated sequences, which are transcribed in a developmentally regulated manner. We anticipate that comprehensive analyses of the expression dynamics, cellular localization, and interaction partners of neural SRS-lncRNAs using the TRE-Ngn2 system (Fig. S3) will illuminate their functional significance in the normal development of the human brain.

## Acknowledgements

We thank Jizhong Zou and Michael Ward for reagents, and Snezhka Oliferenko for valuable discussions.

## Competing interests

The authors have no relevant financial or non-financial interests to disclose.

## Funding

This work was supported by the Biotechnology and Biological Sciences Research Council (grant numbers BB/R001049/1 and BB/V006258/1).

## Author contributions

All authors contributed to the study conception and design. Tek Hong Chung performed the bioinformatics analyses, TRE-Ngn2 differentiation experiments, and RT-qPCR assays. Anna Zhuravskaya generated and characterized induced pluripotent stem cells encoding the TRE-Ngn2 transgene. Eugene Makeyev supervised the project and obtained the funding. The first draft of the manuscript, including the text, figures and the tables, was prepared by Tek Hong Chung and Eugene Makeyev. All authors read and approved the final manuscript.

---

**Figure S1.**
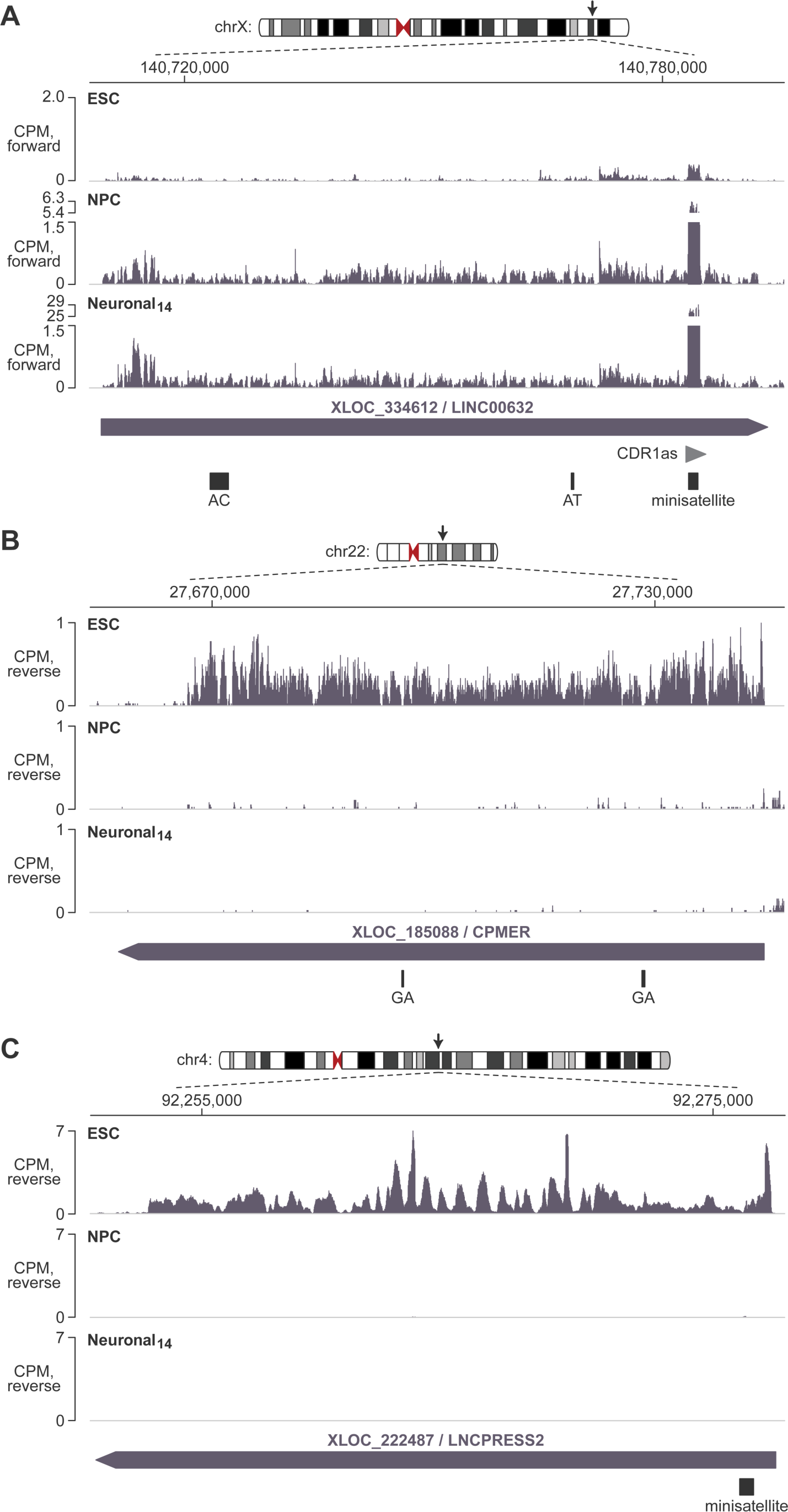
Examples of previously characterized SRS-lncRNAs shortlisted by our workflow. Counts-per-million (CPM) normalized RNA-seq coverage plots are presented for the nuclear fraction of ESC, NPC, and Neuronal_14_ samples for **(A)** XLOC_334612/LINC00632 and CDR1as/CiRS-7, **(B)** XLOC_185088/CPMER, and **(C)** XLOC_222487/LNCPRESS2. Consistent with earlier studies (Barrett et al. 2017; Hansen et al. 2013; Jain et al. 2016; Lyu et al. 2022; Memczak et al. 2013), the SRS-lncRNAs in (A) are upregulated, while those in (B-C) are downregulated during neuronal differentiation. The arrows on the top indicate the chromosomal positions of the three SRS-lncRNA loci. The corresponding SRSs are annotated at the bottom of each panel.

**Figure S2.**
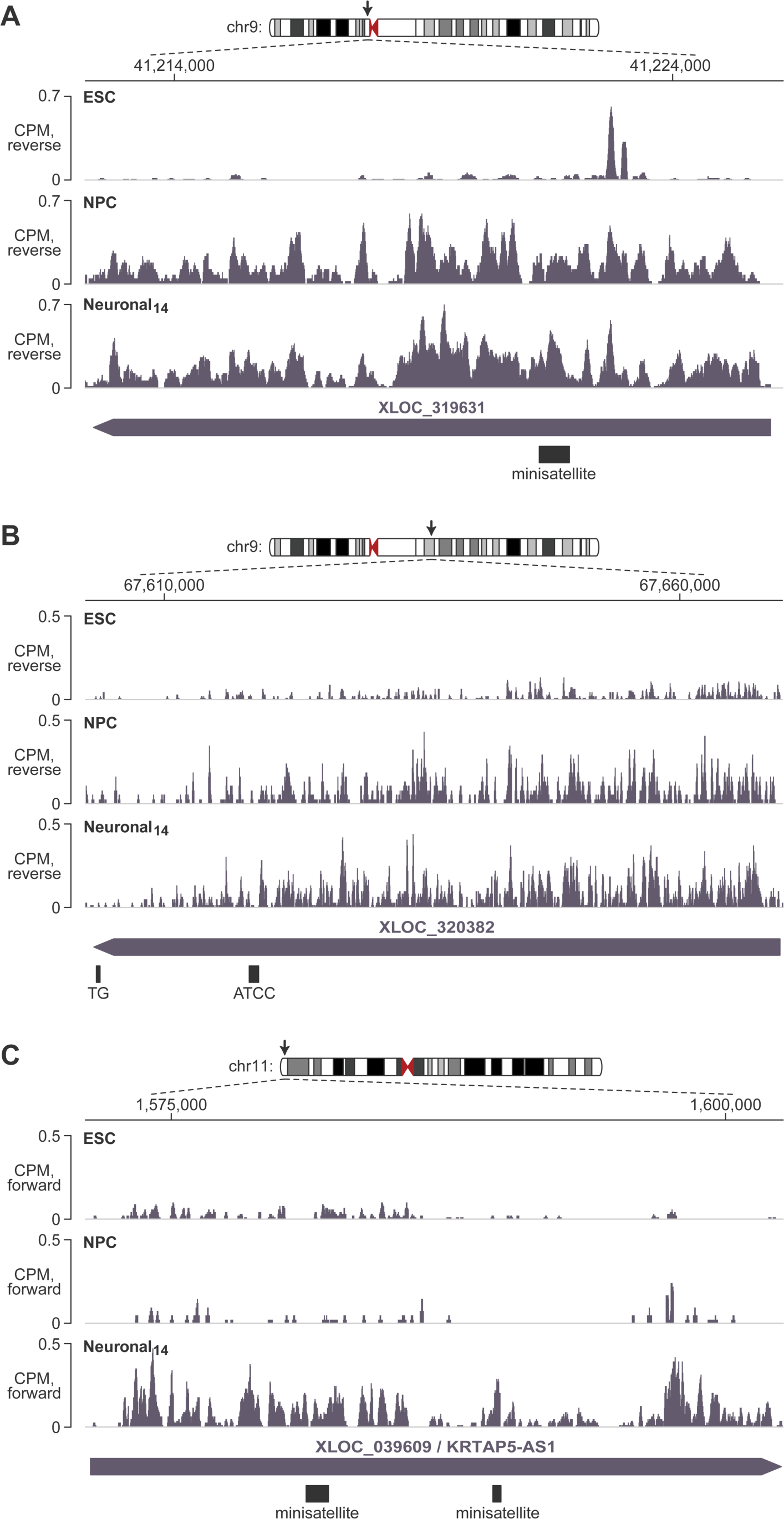
Previously uncharacterized examples of neurally upregulated SRS-lncRNAs. **(A-B)** Shown are CPM-normalized RNA-seq coverage plots for the nuclear fraction of ESC, NPC, and Neuronal_14_ samples, illustrating the SRS-lncRNA XLOC_319631 and XLOC_320382 identified by our pipeline. **(C)** Similar plots for the SRS-lncRNA XLOC_039609/KRTAP5-AS1, previously proposed to function as a competing endogenous RNA in gastric cancer (Song et al. 2017), and shortlisted by our pipeline as neurally upregulated. The arrows on the top mark chromosomal positions of the three SRS-lncRNA loci. The corresponding SRSs are annotated at the bottom of each panel.

**Figure S3.**
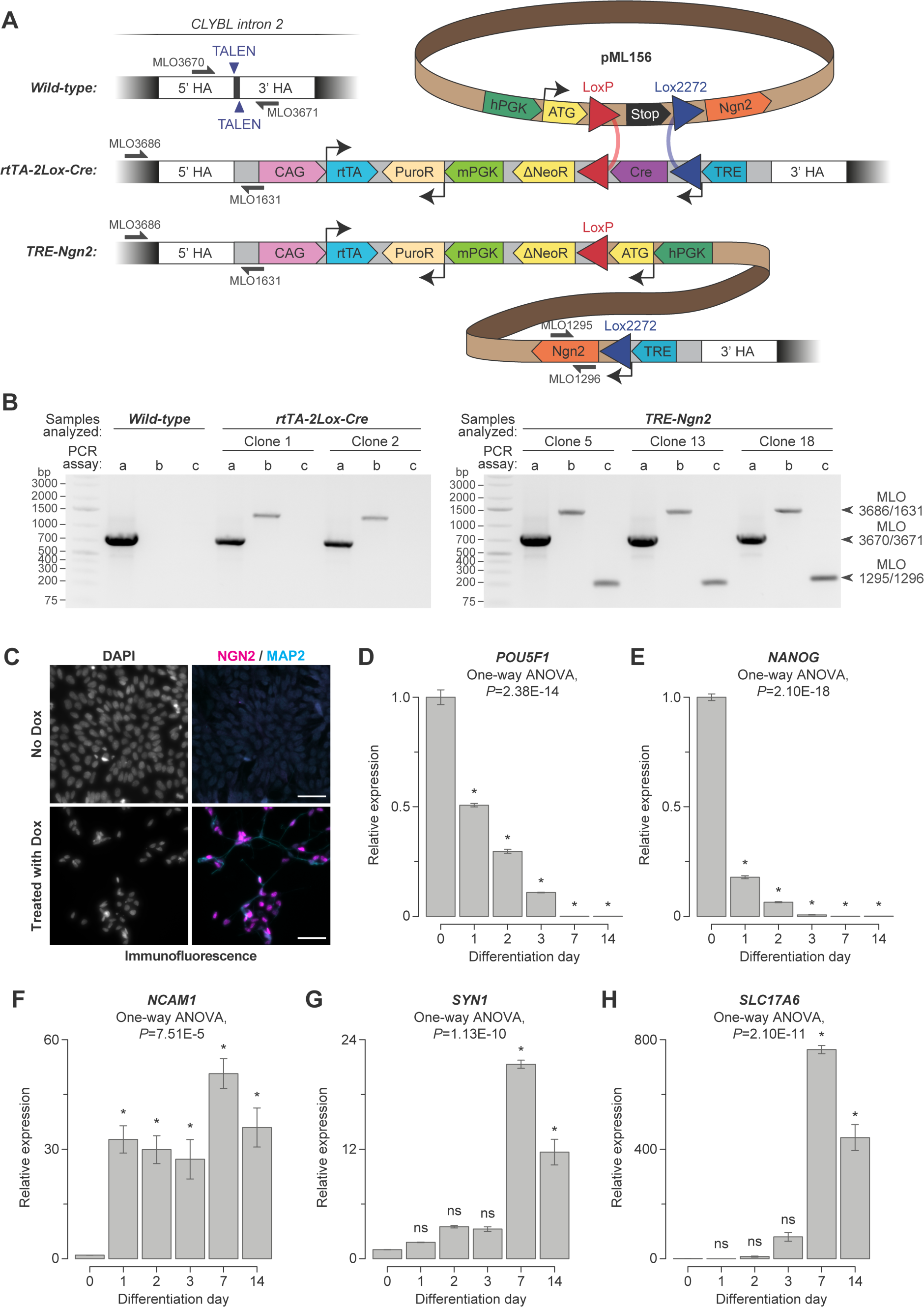
A doxycycline-inducible system for neuronal differentiation of human iPSCs. **(A)** The two-step knock-in approach used in this study to generate TRE-Ngn2 iPSCs. The diagram also indicates the annealing positions of the PCR genotyping primers used to analyze the wild-type *CLYBL* allele (MLO3670-MLO3671, PCR assay a); the knock-in of the rtTA-2Lox-Cre cassette (MLO3686-MLO1631, PCR assay b); and the integration of the Ngn2 transgene (MLO1295-MLO1296, PCR assay c). **(B)** PCR genotyping results for the wild-type iPSCs and their rtTA-2Lox-Cre and TRE-Ngn2 derivatives. The corresponding amplicons are marked on the right. **(C)** An immunofluorescence analysis reveals the expression of transgenic Ngn2 (magenta) and the neuronal marker MAP2 (cyan) in TRE-Ngn2 iPSCs incubated with Dox for 48 hours but not in a control culture grown without Dox. DAPI was used as a nuclear stain. Scale bars, 50 µm. **(D-H)** RT-qPCR analyses of differentiating TRE-Ngn2 cultures for the expression of (D-E) pluripotency markers *POU5F1/OCT4* and *NANOG*; (F) the early neuronal marker *NCAM1*; (G) the mature neuronal maker *SYN1*; and (H) the glutamatergic neuron-specific marker *SLC17A6*/*VGLUT2*. Data are averaged from differentiation experiments using three distinct TRE-Ngn2 iPSC clones ±SEM and compared by one-way ANOVA with Tukey’s post-hoc test. *, Tukey’s *P*<0.05. ns, Tukey’s *P*≥0.05.

## Notes

### Competing Interest Statement

The authors have declared no competing interest.

